# Proneural – Mesenchymal antagonism dominates the patterns of phenotypic heterogeneity in Glioblastoma

**DOI:** 10.1101/2023.11.27.568853

**Authors:** Harshavardhan BV, Mohit Kumar Jolly

**Affiliations:** IISc Mathematics Inititative, Indian Institute of Science, Bengaluru, 560012, India; Department of Bioengineering, Indian Institute of Science, Bengaluru, 560012, India

## Abstract

The aggressive nature of glioblastoma (GBM) – one of the deadliest forms of brain tumours – is majorly attributed to underlying phenotypic heterogeneity. Early attempts to classify this heterogeneity at a transcriptomic level in TCGA GBM cohort proposed the existence of four distinct molecular subtypes: Proneural, Neural, Classical and Mesenchymal. Further, a single-cell RNA-seq analysis of primary tumours also reported similar 4 subtypes mimicking neuro-developmental lineages. However, it remains unclear whether these 4 subtypes identified via bulk and single-cell transcriptomics are mutually exclusive or not. Here, we perform pairwise correlations among individual genes and gene signatures corresponding to these proposed subtypes, and show that the subtypes are not distinctly mutually antagonistic in either TCGA or single-cell RNA-sequencing data. We observed that the proneural (or neural progenitor-like) – mesenchymal axis is the most prominent antagonistic pair, with the other two subtypes lying on this spectrum. These results are reinforced through a meta-analysis of over 100 single-cell and bulk transcriptomic datasets as well as in terms of functional association with metabolic switching, cell cycle and immune evasion pathways. These results suggest rethinking GBM phenotypic characterization for more effective therapeutic targeting efforts.

## 2 Introduction

Glioblastoma Multiforme (GBM) is a primary brain tumour that originates from the glial cells in the brain. It is classified as a grade IV glioma by the World Health Organization (WHO) due to its highly malignant nature (Louis et al., 2021). GBM is not only one of the most common primary brain tumours but also one of the deadliest, with a median survival of only 15 months and a grim 5-year survival of just 6.9% for those diagnosed with it (Chang et al., 2016; Ostrom et al., 2023). The standard of care for GBM patients is surgical resection followed by radiation and temozolomide (TMZ) treatment (Davis, 2016). However, given the aggressive nature of the disease, GBM recurrence is unfortunately very common (Ringel et al., 2015). Recurrent GBMs have an even worse prognosis with a median survival of only 25-40 weeks (Chang et al., 2016). Multiple reasons contribute to the failure of existing treatments: the unique nature of the blood-brain barrier, immunosuppressive environment, highly invasive nature, de novo and acquired drug resistance and phenotypic plasticity and heterogeneity (Dymova et al., 2021; Yabo et al., 2022).

Heterogeneity in GBM exists at many levels: functional, molecular, inter-patient and intra-tumoural (Dymova et al., 2021). It is a major clinical challenge due to varied susceptibilities of cellular subpopulations to treatment. Additionally, GBM cells exhibit a remarkable degree of plasticity, allowing them to adapt and evolve rapidly in response to diverse treatments (Yabo et al., 2022). Similar to observations in breast cancer, lung cancer and melanoma (Goyal et al., 2023; Oren et al., 2021; Ramirez et al., 2016; Sahoo et al., 2021b; Su et al., 2017; Subhadarshini et al., 2023), GBM subpopulations can also undergo genetic and/or non-genetic (reversible) cell-state transitions (Goyal et al., 2023; Stepanenko et al., 2016; L. Wang et al., 2022), often driving drug resistance and eventual GBM recurrence. Understanding the molecular underpinnings of such plasticity and heterogeneity is, therefore, essential.

An earlier key study characterising heterogeneity in GBM used transcriptomic data from the TCGA-GBM cohort and reported 4 subtypes: Proneural (TCGA-PN), Neural (TCGA-NL), Classical (TCGA-CL), and Mesenchymal (TCGA-MES) (Verhaak et al., 2010). Except for the neural subtype, other subtypes were associated with specific gene abnormalities: EGFR alterations in classical, NF1 mutations in mesenchymal, and PDGFRA and IDH1 mutations in proneural. Upon quantifying the enrichment of gene expression profiles of neural cell types in GBM, the proneural samples were highly enriched for oligodendrocytic signature, while the mesenchymal ones were strongly associated with cultured astroglial signature. Further analysis suggested that the neural subtype may be associated with contamination of normal neural cells in the GBM sample, indicating that proneural, mesenchymal and classical phenotypes may be the GBM-specific ones (Gill et al., 2014; Q. Wang et al., 2017). Multi-region tumour sampling and single-cell RNA-sequencing (scRNA-seq) demonstrated that these different molecular subtypes can co-exist in the same tumour specimen (Gill et al., 2014; Patel et al., 2014; Sottoriva et al., 2013). These observations are reminiscent of intra-tumour heterogeneity along the proliferative-invasive spectrum in melanoma samples (Pillai et al., 2022) and along the epithelial-mesenchymal axis in carcinomas (Bocci et al., 2019; Brown et al., 2022; Malagoli Tagliazucchi et al., 2023).

To further characterise intra-tumour heterogeneity, scRNA-seq of GBM tumours from 28 adult and pediatric patients were used, and 4 meta-modules mimicking neuro-developmental lineages were identified: neural-progenitor-like (NPC-like), oligodendrocyte-progenitor-like (OPC-like), astrocyte-like (AC-like), and mesenchymal-like (MES-like) states. Each tumour analysed contained cells in at least 2 of these 4 cell-states, with varied relative ratios. These subtypes had amplifications of CDK4, PDGFRA, EGFR and NF1 respectively. Consistently, AC-like and MES-like states were found to correspond to TCGA-CL and TCGA-MES subtypes, respectively, while TCGA-PN subtype corresponded to a combination of OPC-like and NPC-like cell-states (Neftel et al., 2019). Together, these landmark studies in GBM heterogeneity posited the (co-) existence of four distinct cellular states. However, whether these four cell-states are mutually exclusive or antagonistic among one another or not remains elusive.

Here, we first quantify pairwise correlations among the scores of all four proposed subtypes in TCGA and scRNA-seq datasets based on which these subtypes were proposed and demonstrate that only proneural (or neural-progenitor-like) - mesenchymal pair shows a strong antagonistic pattern. Next, we show that this antagonism is also present at the individual gene level among these two states. Finally, a meta-analysis of more than 100 bulk and single-cell RNA-seq GBM datasets endorses this analysis and reveals functional differences between proneural and mesenchymal states in terms of their association with cell cycle, metabolic status and immune evasion. Our results thus suggest that the proneural-mesenchymal switch dominates the patterns of GBM phenotypic heterogeneity.

## 3 Results

### 3.1 Scoring of subtype-specific signatures reveal that not all cell-states are mutually antagonistic

The four proposed sub-types were classified at a transcriptomic level and thus characterised by sets of genes highly expressed in each of them (Neftel et al., 2019; Verhaak et al., 2010). Thus, we quantified the enrichment of different gene sets in each sample in a given cohort. If the proposed subtypes correspond to distinct cell-states, we expect to see antagonism or independence in the expression patterns of these gene sets. Mutually exclusive or antagonistic gene expression profiles can emerge from one set of genes/master regulators of one phenotype suppressing those from the other sets, as witnessed in other cancers exhibiting phenotypic plasticity (Hoek et al., 2008; Zhang et al., 2018). Thus, antagonism in gene regulatory networks is expected to manifest as a negative correlation between the enrichment scores of such gene sets. However, if the gene modules corresponding to different phenotypes do not regulate one another, we expect to see no correlation among their scores. As expected, the genesets corresponding to these four subtypes - (TCGA-CL, TCGA-MES, TCGA-PN, TCGA-NL) (Verhaak et al., 2010) or (NPC-like, APC-like, OC-like and MES-like) (Neftel et al., 2019) - do not show much overlap (Figure 1A).

**Figure 1:**
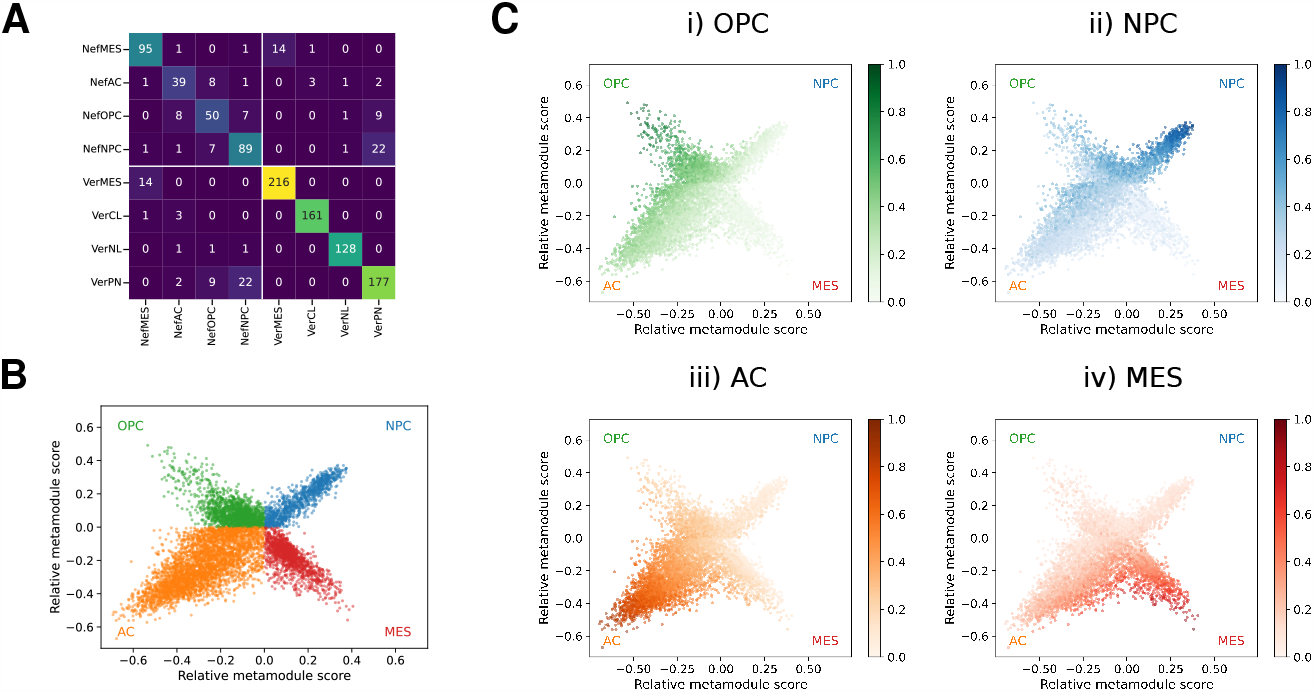
Subtypes defined for GBM: **A**: Representation of number of genes within each geneset, highlighting the common genes between different genesets. **B**: Recreation of the two-dimensional representation of cellular states as defined in Neftel et al., 2019. **C**: Two-dimensional representation coloured by scores for each subtype: (i) OPC-like, (ii) NPC-like, (iii) AC-like, (iv) MES-like.

We first recreated the graphical representation presented earlier (Neftel et al., 2019) using corresponding enrichment scores of the signatures corresponding to the proposed four subtypes (Figure 1B). This projection adopts a discontinuous x-axis, which, while being a valid stylistic choice, introduces a visual separation that may inadvertently emphasise the distinctions between cell states, leading to the impression that the four states are all mutually antagonistic and that they correspond to four independent dimensions. Moreover, the utilisation of the “max” operation between sets of subtypes for the y-axis could contribute to this exaggeration. Thus, we questioned whether this portrayal could be inadvertently obscuring the true and intricate relationships between the identified states. First, we quantified the scores of individual subtypes on this projection (Figure 1C). The scores of a particular subtype are relatively high in the “arm” corresponding to that subtype but not exclusive; also, the discontinuity of the x-axis becomes clearly visible with a break in the gradient. This analysis suggested that the use of a discontinuous x-axis was at least partly distorting the true relationship among the proposed subtypes in the underlying gene expression space, and thus, unlike the visual impression created, the four subtypes may not be as mutually exclusive among one another.

We investigated the two source datasets: GSE131928 and TCGA-GBM, from which the Neftel signature and Verhaak signature were derived, respectively. In both these groups of four genesets each, we noticed that not all pairwise correlations are negative; instead, many pairs show positive correlation as well, indicating that they are not distinct (Figure 2A). Particularly, we noticed that the Neftel NPC-like and MES-like signatures are negatively correlated and, hence, antagonistic. Similarly, for the Verhaak signatures, the PN and MES pair are antagonistic. When comparing across the Neftel and Verhaak signatures, we had the following observations: a) Neftel NPC-like and Verhaak PN are positively correlated with each other, b) Neftel MES-like and Verhaak MES are positively correlated with one another, and c) both Neftel NPC and Verhaak PN are negatively correlated with the MES signatures from both the geneset cohorts, further endorsing previous observations about NPC-like and PN may correspond to similar cell-states (Neftel et al., 2019).

**Figure 2:**
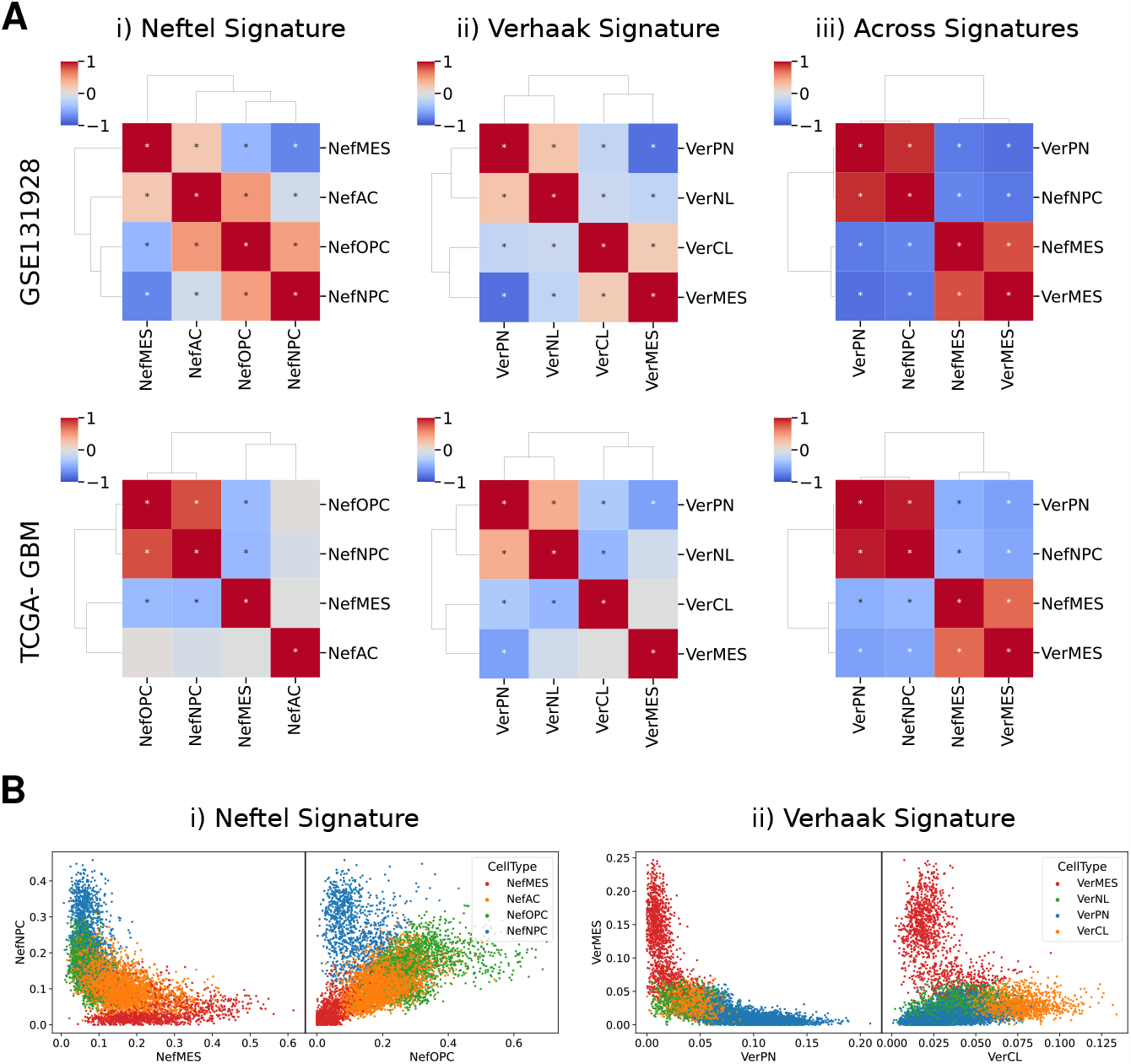
Trends revealed by the Signature scores: **A**: Heatmaps visualizing spearman correlation of ssGSEA and AUCell scores for subtypes in the (i) Neftel signature, (ii) Verhaak signature, and (iii) comparison between subtypes across these signatures across (top) GSE131928 and (bottom) TCGA-GBM datasets. The presence of “*” indicates that the correlation has a p-value < 0.05. **B**: Scatterplots showing AUCell scores from the GSE131928 dataset, comparing scores between genesets that exhibit negative and positive correlations. For the (i) Neftel signature, the comparison is between NPC-like-vs-MES-like and NPC-like-vs-OPC-like, and for the (ii) Verhaak signature, the comparison is between MES-vs-PN and MES-vs-CL.

The difference between the pairs of states that are negatively and positively correlated is further illustrated through the scatter between the pairs of scores coloured by the maximum score in the GSE131928 dataset (Figure 2B). The cells with high PN/NPC enrichment tend to have very low or no expression of MES genes. Interestingly, the cells with high enrichment of AC-like or OPC-like in Neftel signatures and CL or NL in Verhaak signatures lie in between the gradient of scores in these axes. On the other hand, the cells that are highly enriched for OPC-like tend to have comparable expression of NPC-like as well. This trend is observed for the other pairs as well (Figure S1A) and in the TCGA-GBM dataset (Figure S1B).

Together, our results indicate that across the two datasets used for identifying the two sets of four subtypes (Neftel et al., 2019; Verhaak et al., 2010), the proneural (PN) (or equivalently NPC-like) and mesenchymal (or equivalently MES-like) show strongest antagonistic trends, despite little overlap in gene signatures of PN & NPC-like or MES & MES-like.

### 3.2 NPC/PN-MES antagonism is prevalent at the gene level as well

After investigating pairwise correlations at the gene signature level, we then focused on the gene level to confirm that our observations were not an artifact of the gene set enrichment scoring methods. Thus, we calculated pairwise correlation among all genes involved in the 4 Neftel gene sets together and noticed that most MES genes correlated positively with one another, most NPC genes correlated positively with each other, but most MES genes were antagonistic to most NPC ones (Figure 3A). Similar observations were made for Verhaak signatures as well – where genes from PN, MES and CL clustered separately - and for both the source datasets – GSE 131928 and TCGA-GBM (Figure 3A, Figure S2A). These observations suggest that antagonism seen among the enrichment scores of MES vs NPC or PN emerges from coordinated expression profiles of genes showing a negative correlation. To quantify these associations, we define J-metric (Equation 2) that quantifies the extent of positive correlation among genes in the same geneset and extent of negative correlation among those across genesets (Chauhan et al., 2021). J-metric calculations further support our observations that the MES-NPC or the MES-PN gene sets are the most antagonistic ones among all six pairwise correlations (Figure 3B,Figure S2B) (Neftel: MES-AC, MES-OPC, MES-NPC, AC-OPC, AC-NPC, OPC-NPC and Verhaak: MES-CL, MES-NL, MES-PN, CL-NL, CL-PN, NL-PN).

**Figure 3:**
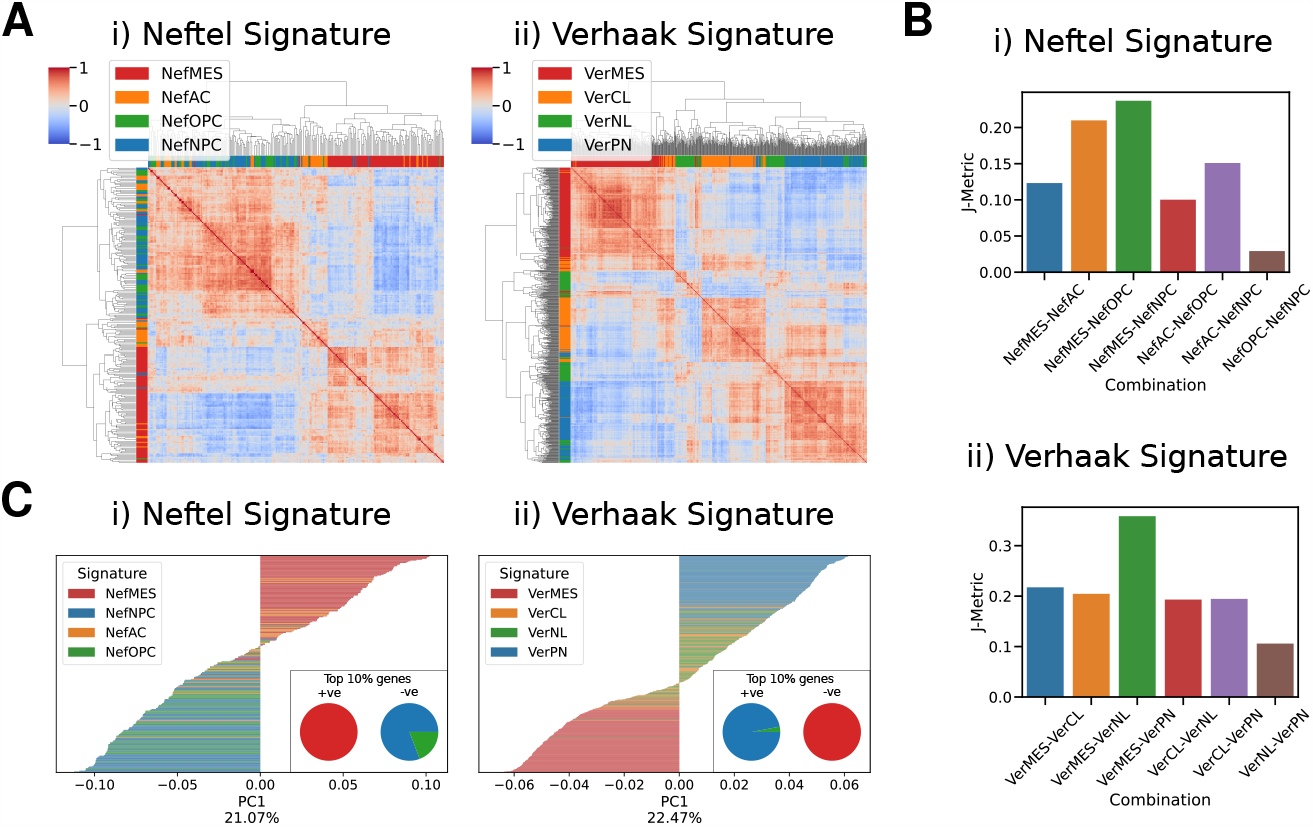
Analysis for antagonism of signature genes in the TCGA-GBM dataset: **A** Heatmaps visualizing spearman correlation of gene expression levels in the (i) Neftel signature and (ii) Verhaak signature. The colors beside the heatmap represent the geneset to which each gene belongs. **B**: J-Metric values for various combinations of subtypes for (i) Neftel signature and (ii) Verhaak signature. **C**: PC1 loadings of PCA based on the gene expression levels in the (i) Neftel signature and (ii) Verhaak signature. The inset shows the geneset to which the top 10% of genes belong for both positive and negative loadings.

Next, we interrogated whether this antagonistic relationship also corresponds to the highest variance. To investigate this, we conducted Principal Component Analysis (PCA) on the gene expression levels within each signature. The first principal component (PC1) represents the axis that captures the greatest variance. Looking at the PC1 loadings for TCGA and GSE131928 (Figure 3C, Figure S2C), we noticed that for the Neftel signatures, MES genes dominate on loadings with positive PC1 coefficient and NPC genes on the negative loadings. Similarly, for the Verhaak signatures, the PN genes dominate on the positive loadings and the MES genes on the negative loadings. Together, these results suggest that the MES-NPC/PN antagonism seen at signature levels percolates to individual genes, and among all the six possible pairwise analyses, MES and NPC (or PN) genes show distinct mutually exclusive patterns.

### 3.3 Meta-analysis across multiple transcriptomic datasets reveal the functional consequences of NPC/PN-MES antagonism

To test for the generalisability of our results, we investigated if the trends we observe hold up across multiple transcriptomic datasets. We examine 80 bulk RNA-sequencing datasets with samples spanning across patient tumour samples, mouse models and cell-lines to capture the maximum variability. For datasets with samples across different organisms (mouse models, patient samples) and contexts (in vitro, in vivo), we segregated them accordingly and combined the samples, totalling to 90 consolidated samples across all datasets. In each consolidated sample, we calculated the correlation coefficient between the scores of each pair of subtypes. For the correlations between Neftel MES-like and Neftel NPC-like, out of the 38 samples that show a significant correlation, 33 samples (86.8 %) are negatively correlated, with just 5 being positive. This pair shows the most bias for being negatively correlated among all the possible combinations, with a few of the other pairs showing bias towards being positively correlated (Figure 4A). Similarly, for the correlations between Verhaak MES and Verhaak PN, of the 48 samples that show a significant correlation, 45 (93.8 %) of them are negatively correlated, with just 3 being positive. This pair shows the most bias for negatively correlated among all possible pairwise comparisons.

**Figure 4:**
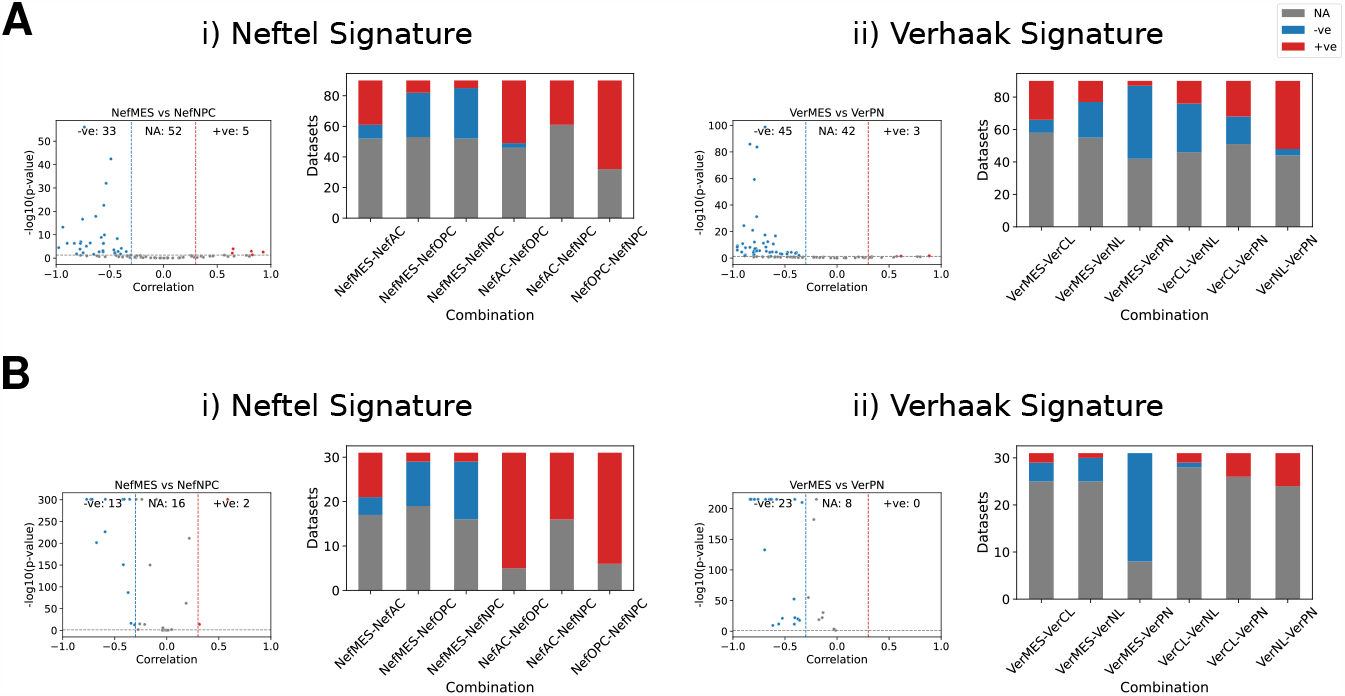
Meta-analysis for trends in correlation of subtypes: **A**: Analysis of Bulk RNASeq datasets. **B**: Analysis of Single-Cell RNASeq datasets. The left panel displays the volcano plot of correlation between NPC-vs-MES and PN-vs-MES for (i) Neftel Signature and (ii) Verhaak Signature. A dataset with correlation coefficient < -0.3 and p-value < 0.05 is categorised as negatively correlated, with correlation coefficient *>* 0.3 and p-value < 0.05 is categorised as positively correlated and NA otherwise. The right panel displays the stacked bar plot of these counts for correlation between different combinations of subtypes.

However, when analysing bulk transcriptomic datasets, positive correlations could also arise due to the co-occurrence of those subtypes in a tumour and thus may not reflect their genuine transcriptomic associations. To address this, we have looked at 26 single-cell RNASeq datasets, with 30 consolidated samples as well. Again, here we see the Neftel MES and Neftel NPC pair to have the most bias for negatively correlated of 13 (86.7%) out of 15 significantly correlated, which is the highest bias for negatively correlated (Figure 4B). For the Verhaak signature, the MES-PN pair have the most bias for negatively correlated of 23 (100%) out of 23 significantly correlated.

Next, we checked if the antagonism between MES-NPC/PN has any functional relevance. To characterise the immune infiltration, we looked at the correlation of PD-L1 signature scores with the subtype specific scores. With the bulk datasets, Neftel NPC and Verhaak PN showed the highest negative correlation with PD-L1 of 33 out of 34 (97%) and 35 out of 38 (92.1%) respectively. Meanwhile, the Neftel and Verhaak MES showed the highest positive correlation with PD-L1 of 49 out of 55 (89.1%) and 55 out of 60 (91.7%) respectively (Figure 5A). To characterise the metabolic activity, we looked at the correlation of Glycolysis signature scores with the subtype specific scores. With the bulk datasets, Neftel NPC and Verhaak PN showed the highest negative correlation of 24 out of 39 (61.5%) and 29 out of 42 (69%) respectively, while, the Neftel and Verhaak MES showed the highest positive correlation of 60 out of 61 (98.4%) and 57 out of 60 (95%) respectively (Figure 5B). Similarly, for proliferative activity, we looked at the correlation with KEGG cell cycle signature score. With the bulk datasets, Neftel NPC and Verhaak PN showed the highest positive correlation of 18 out of 28 (64.3%) and 34 out of 40 (85%) respectively, while, the Neftel and Verhaak MES showed the highest negative correlation of 19 out of 38 (50%) and 24 out of 60 (63%) respectively (Figure S3A).

**Figure 5:**
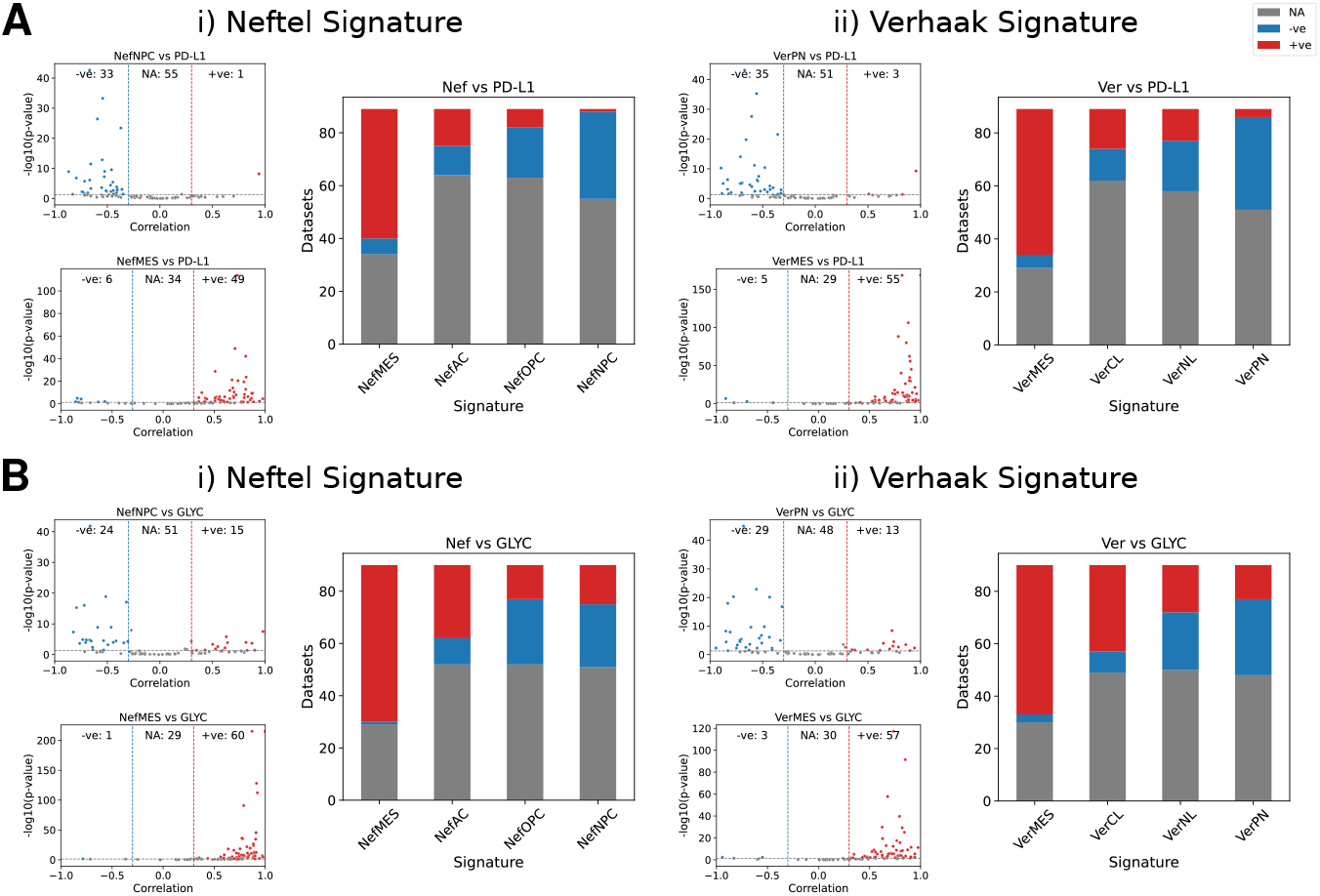
Meta-analysis for trends in correlation with immune and metabolic signatures for bulk datasets: **A**: Correlation of subtypes with PD-L1 Signature score. **B**: Correlation of subtypes with Hallmark Glycolysis score. The left panel displays the volcano plot of correlation between (top left) Signature-vs-NPC/PN and (bottom left) Signature-vs-MES for (i) Neftel Signature and (ii) Verhaak Signature. A dataset with correlation coefficient < -0.3 and p-value < 0.05 is categorised as negatively correlated, with correlation coefficient *>* 0.3 and p-value < 0.05 is categorised as positively correlated and NA otherwise. The right panel displays the stacked bar plot of these counts for correlation between different subtypes with either of the signatures.

Similar trends in general are observed with the single cell datasets as well, however, these should be treated with caution given the low number (<10) of cases showing significant correlation. Generally, the MES signatures are more negatively correlated with KEGG cell cycle (Figure S3B), and, positively with PD-L1 (Figure S3C) and glycolyis (Figure S3D) while, NPC/PN are positively with KEGG cell cycle and negatively with PD-L1 and glycolysis. Together these results points towards NPC/PN and MES occupying opposing ends of the functional spectrum.

## 4 Discussion

Our in-depth analysis underscores that not all four proposed GBM subtypes can be unequivocally categorized as distinct entities. It is apparent that, within this classification, some subtypes exhibit a level of similarity that prevents us from asserting their distinctiveness. However, we noticed consistently that the proneural/mesenchymal axis (as per Verhaak et al., 2010 classification) or neural progenitor-like/mesenchymal-like axis (as per Neftel et al., 2019 classification) are the most notably antagonistic pairs. This antagonistic relationship, together with their inverse relationships with functional attributes (cell cycle, immune evasion, metabolism) as identified via our meta-analysis, implies a clear demarcation between these two subtypes, rendering them genuinely distinct states.

Our results are reminiscent of previous single-cell/single-nucleus RNA-seq observations of a cohort of primary tumours showing that phenotypes of proliferating GBM cells lie along a single axis of variation ranging from proneural to mesenchymal (L. Wang et al., 2019). Diverse biological phenotypes being mostly explained by Principal Component 1 (PC1) axis has also been observed for other instances of cancer cell plasticity: epithelial-mesenchymal transition in carcinomas, and proliferative-invasive switch in melanomas (Hari et al., 2023), suggesting this feature to be a more generic occurrence than only in GBM. Transcriptional variability along the proneural-mesenchymal axis also impacts drug sensitivity (Meyer et al., 2015), similar to the association of EMT with resistance to chemotherapy, targeted therapy and immune checkpoint blockade therapy (Dudás et al., 2020; Gu et al., 2023; Sahoo et al., 2021b). Consistently, recurrent GBMs are enriched in a mesenchymal state (L. Wang et al., 2022). This variability also has implication in the development of targeted therapy. For example, a drug aimed at the NPC-like state can also impact cells in the OPC-like state, given their transcriptional similarity. Conversely, the MES-like state is expected to be refractory to the drug targeting NPC-like state.

Our meta-analysis suggests that mesenchymal subtype is relatively enriched in glycolytic traits, a behaviour validated in metabolic differences between proneural and mesenchymal tumor-initiating cells (Mao et al., 2013; Seliger et al., 2022). The observed association of PD-L1 with mesenchymal subgroup in GBM (Nduom et al., 2015; Ricklefs et al., 2018; Wu et al., 2022) also reinforces our analysis; functionally, PD-L1 can trigger more aggressive GBM behaviour through the downstream Ras/Erk signaling. The independent association of both PD-L1 and glycolysis with worse patient outcomes in GBM (Nduom et al., 2015; Z. Wang et al., 2016) suggests more aggressive behaviour of MES phenotype, given the enrichment of both glycolysis and PD-L1 levels in MES subtype. Concurrent enrichment of glycolysis and PD-L1 was found to be associated with worse overall survival across multiple cancer types as well (Muralidharan et al., 2022).

Our results establish important functional differences between proneural and mesenchymal states and suggest rethinking GBM phenotypic classification. Thus, future efforts to characterize GBM phenotypic plasticity and heterogeneity should incorporate both genetic and non-genetic (transcriptional, epigenetic, metabolic) components, given increasing evidence about the interplay of both components in cancer cell adaptation to therapeutic attacks (Fernandez-Mateos et al., 2023; Garofano et al., 2021; Salgia & Kulkarni, 2018). Further, the mapping of underlying gene regulatory networks enabling phenotypic heterogeneity – as done in EMT, neuroendocrine differentiation in small cell lung cancer, and phenotypic switching in melanoma (Ozen & Lopez, 2023; Pillai & Jolly, 2021; Silveira et al., 2020; Steinway et al., 2015; Udyavar et al., 2017) - is still in its infancy in GBM (Larsson et al., 2021; Perez-Aliacar et al., 2023). A hallmark of these networks across the cancer types is the presence of “teams” of mutually inhibitory players that can allow for co-existence of cell-states in a population and most variability in phenotypic space residing along the PC1 axis (Chauhan et al., 2021; Hari et al., 2022, 2023; Pillai & Jolly, 2021). Whether such “teams” exist for proneural vs mesenchymal phenotypes remains elusive as of yet.

Overall, the results presented here across bulk and single-cell transcriptomics demonstrate that proneural-mesenchymal axis dominates the patterns of phenotypic heterogeneity in GBM, and cells lying along this axis have varied functional traits in terms of metabolism and immune-evasion. Such extensive meta-analysis conducted for over 100 datasets offers an unbiased view of GBM heterogeneity, and highlights clinical implications of its accurate classification at both diagnostic and therapeutic aspects. Moreover, it can be a first key step to narrow down on the molecular mechanisms shaping these low-dimensional phenotypic heterogeneity patterns.

## 5 Methods

### 5.1 Datasets and preprocessing

All the datasets are publicly available and given in Table S1 and Table S2 for single-cell and bulk RNA sequencing datasets respectively. The datasets with the GSE IDs are available and downloaded from the Gene Expression Omnibus (GEO) website. Other datasets such as The Cancer Genome Atlas (TCGA), Chinese Glioma Genome Atlas (CGGA), Glioma Longitudinal AnalySiS (GLASS), Cancer Cell Line Encyclopedia (CCLE), QCell were downloaded from their respective website.

We consolidated all the samples from the same datasets into a single count matrix, except for instances where there were variations in the model system or sequencing platform, under which circumstances they were treated separately. Only the tumour cells were selected for the single-cell datasets for which metadata was available. Finally, all the bulk datasets, were normalised to log transcripts per million (*log*_2_(*TPM*)) and, for the single cell datasets, they were log normalised.

### 5.2 Scoring of Signatures

The Verhaak signatures and the pathways signatures for Hallmark Glycolysis and KEGG cell-cyle were obtained from MSigDB (Subramanian et al., 2005). The Neftel signatures and signature for PD-L1 were obtained from previous reports (Neftel et al., 2019; Sahoo et al., 2021a). These signatures are given in Table S3.

The scoring of bulk datasets were done using ssGSEA (Barbie et al., 2009) given by:

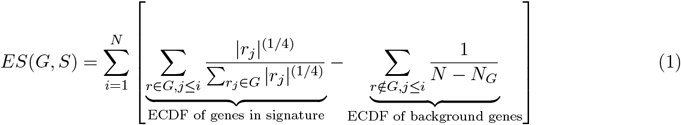

Where,

*G*_*i*_ = Gene Set i

*N*_*i*_ = Number of genes in set i

*r*_*j*_ = Rank of gene j

The implementation of ssGSEA in GSEAPy (Fang et al., 2023) was used and the normalised expression scores were used for further analysis. The scoring of the single-cell datasets were done using AUCell from the SCENIC package in R (Aibar et al., 2017). Pairwise correlations between the scores were done with `spearmanr` function in SciPy.

### 5.3 Correlation of Gene expression

The expression levels of the genes for a particular signature were selected. The pairwise spearman correlations were computed using the cor function in R. These correlations were then used to group together genes with similar expression patterns through hierarchical clustering, which was accomplished using the `clustermap` function in seaborn.

To quantify the antagonism between the different genesets, the J-Metric (Chauhan et al., 2021) was used given by:

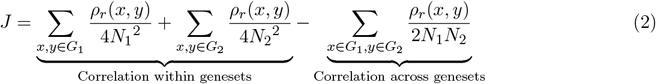

Where,

*G*_*i*_ = Gene Set i

*N*_*i*_ = Number of genes in set i

*ρ*_*r*_(*x, y*) = Spearman correlation of gene x with gene y

A higher value of J-metric corresponds to more antagonism between the genesets. A value closer to 1 indicates that the within geneset correlation is more positive and the across geneset correlation is more negative. Whereas, a value closer to 0 indicates the correlation within geneset and across geneset to be of similar levels.

### 5.4 PCA of Gene expression

Principal Component Analysis (PCA) was done on the expression levels of all the genes for each signature using `prcomp` function in R. Subsequently, the loadings of the first principal component (PC1) were visualized in a bar plot, sorted by their values. To further analyse these results, the top 10% of genes were selected based on their loadings in PC1, both in the positive and negative directions. The counts of genes belonging to each geneset were then quantified.

### 5.5 Code availability

The scripts used for analysis and the processed data are available at: https://github.com/Harshavardhan-BV/GBM4states

## 6 Author contributions

MKJ conceptualised and designed the study. HBV performed the analysis. HBV and MKJ wrote and edited the manuscript.

## 7 Acknowledgements

HBV is supported by the Prime Minister’s Research Fellowship (PMRF). MKJ was supported by Ramanujan Fellowship (SB/S2/RJN-049/2018) by the Science and Engineering Research Board (SERB), Department of Science and Technology, Government of India.

## 8 Conflict of Interest

The authors declare no conflict of interest.

## Supplementary Material

**Figure S1:**
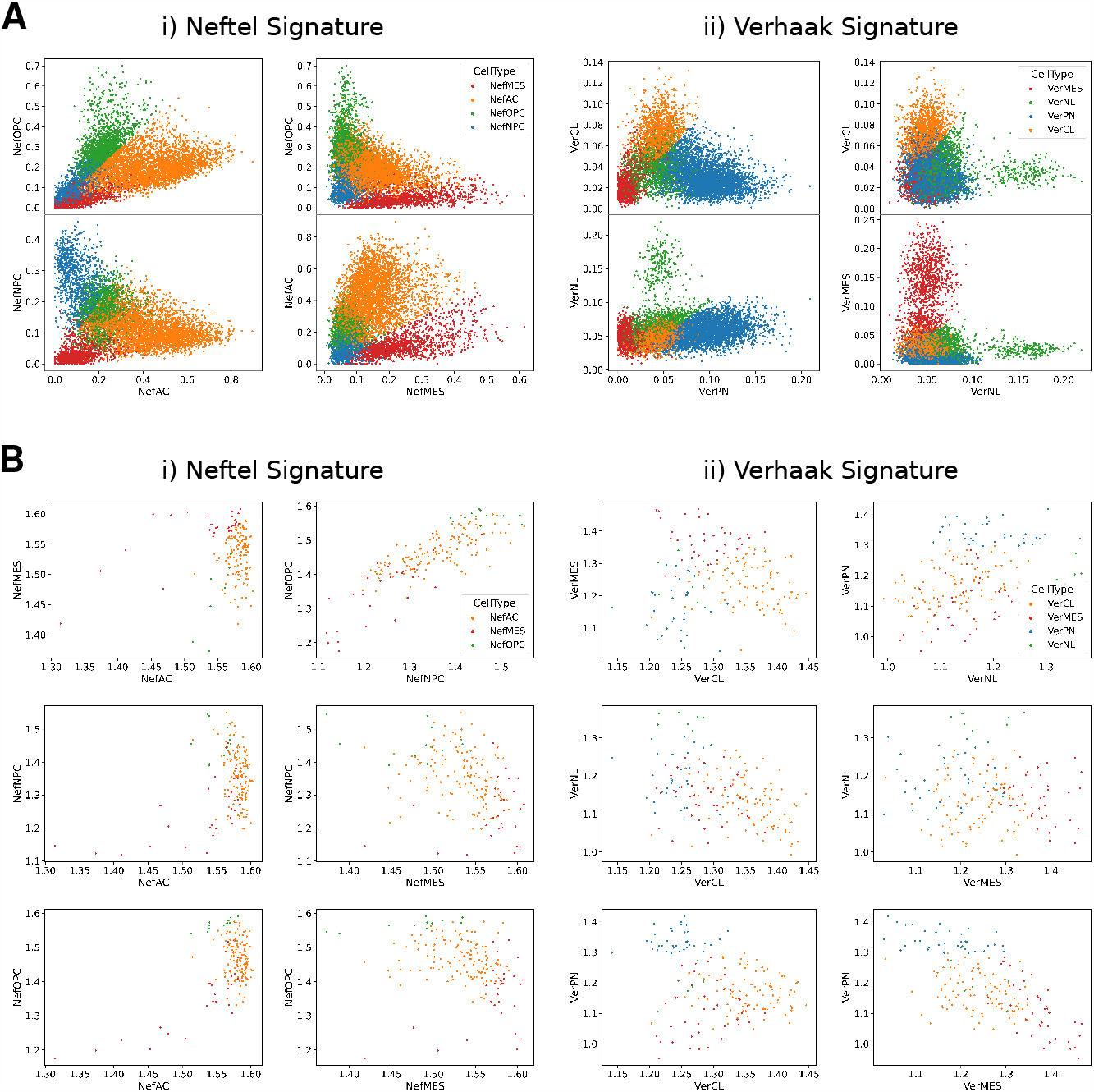
**A**: Scatterplots showing AUCell scores from the GSE131928 dataset for the remaining combinations of genesets for (i) Neftel signatures and (ii) Verhaak signatures. **B**: Scatterplots showing ssGSEA scores from the TCGA dataset for all the combinations of genesets. The left row pertains to Neftel signatures, while the right row pertains to Verhaak signatures.

**Figure S2:**
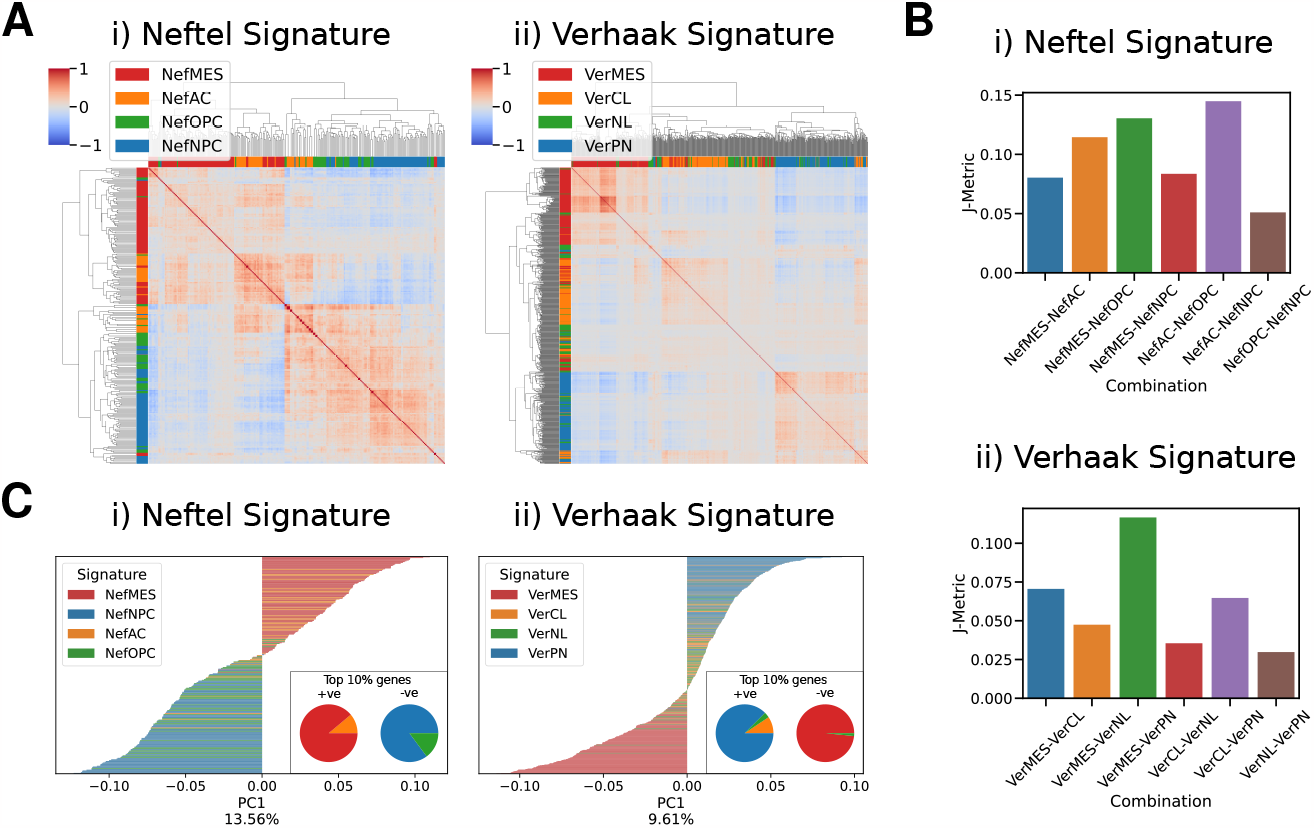
Analysis for antagonism of signature genes in the GSE131928 dataset: **A** Heatmaps visualizing spearman correlation of gene expression levels in the (i) Neftel signature and (ii) Verhaak signature. The colors beside the heatmap represent the geneset to which each gene belongs. **B**: J-Metric values for various combinations of subtypes for (i) Neftel signature and (ii) Verhaak signature signature. **C** PC1 loadings of PCA based on the gene expression levels in the (i) Neftel signature and (ii) Verhaak signature. The inset shows the geneset to which the top 10% of genes belong for both positive and negative loadings.

**Figure S3:**
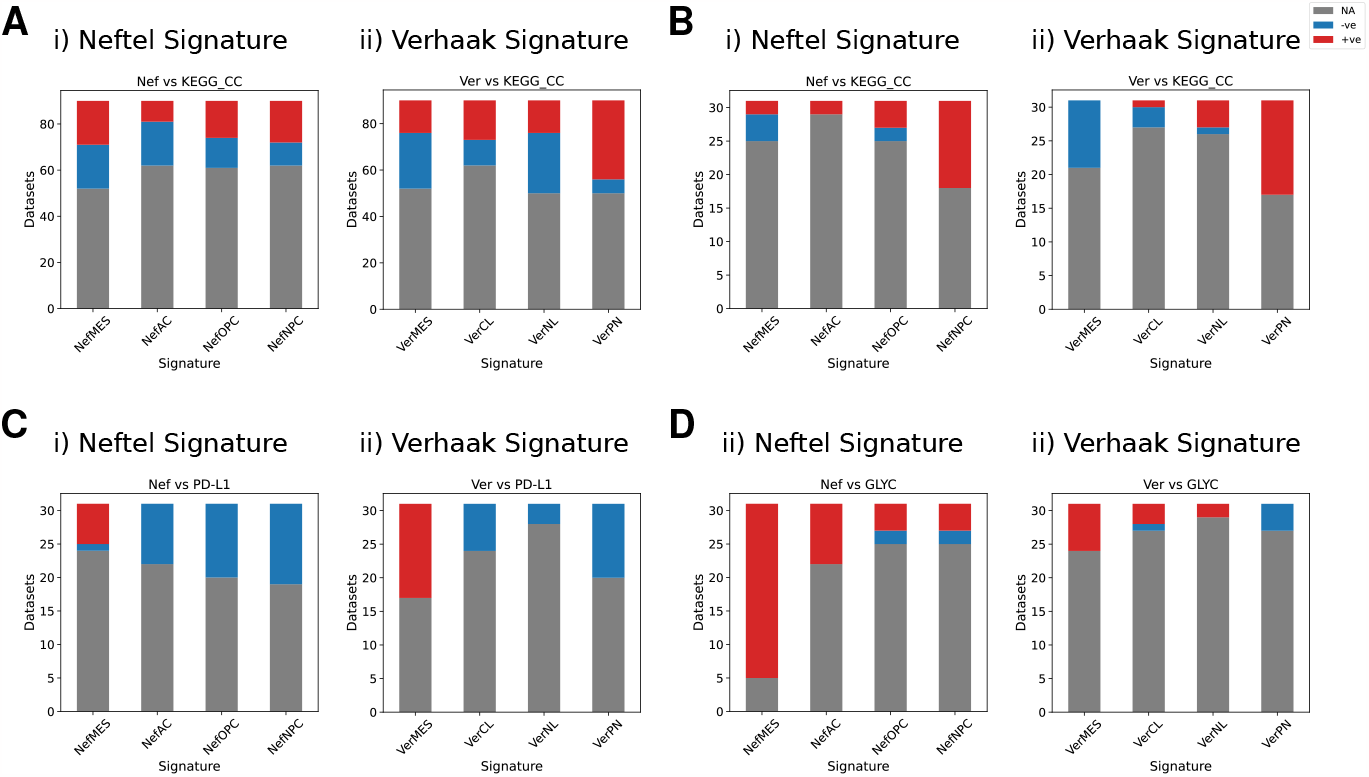
Meta-analysis for trends in correlation with cellcycle, immune and metabolic signatures for bulk and single datasets: **A**: Stacked bar plot illustrating the counts of correlations between subtypes and KEGG Cell Cycle signature score in bulk datasets. **B**: Correlations between subtypes and KEGG Cell Cycle signature score for Single-Cell datasets. **C**: Correlations between subtypes and PD-L1 signature score for Single-Cell datasets. **D**: Correlations between subtypes and Hallmark Glycolysis score for Single-Cell datasets.

## Notes

### Competing Interest Statement

The authors have declared no competing interest.

